# A blood-based signature of cerebrospinal fluid A*β*_1–42_ status

**DOI:** 10.1101/190207

**Authors:** Benjamin Goudey, Bowen J Fung, Christine Schieber, for the Alzheimer’s Disease Metabolomics Consortium, for the Alzheimer’s Disease Neuroimaging Initiative, Noel G Faux

**Author notes:** A complete listing of ADNI investigators can be found at: http://adni.loni.usc.edu/wp-content/uploads/how_to_apply/ADNI_Acknowledgement_List.pdf. A complete listing of ADMC investigators can be found at: https://sites.duke.edu/adnimetab/who-we-are/. These authors contributed equally to this work (Benjamin Goudey, Bowen J Fung). Data used in preparation of this article were generated by the Alzheimer’s Disease Metabolomics Consortium (ADMC). As such, the investigators within the ADMC provided data but did not participate in analysis or writing of this report. Data used in preparation of this article were obtained from the Alzheimer’s Disease Neuroimaging Initiative (ADNI) database (adni.loni.usc.edu). As such, the investigators within the ADNI contributed to the design and implementation of ADNI and/or provided data but did not participate in analysis or writing of this report.

## Abstract

It is increasingly recognized that Alzheimer’s disease (AD) exists before dementia is present and that shifts in amyloid beta occur long before clinical symptoms can be detected. Early detection of these molecular changes is a key aspect for the success of interventions aimed at slowing down rates of cognitive decline. Recent evidence indicates that of the two established methods for measuring amyloid, a decrease in cerebrospinal fluid (CSF) amyloid *β*_1*–*42_ (A*β*_1*–*42_) may be an earlier indicator of Alzheimer’s disease risk than measures of amyloid obtained from Positron Emission Tomography (PET). However, CSF collection is highly invasive and expensive. In contrast, blood collection is routinely performed, minimally invasive and cheap. In this work, we develop a blood-based signature that can provide a cheap and minimally invasive estimation of an individual’s CSF amyloid status using a machine learning approach. We show that a Random Forest model derived from plasma analytes can accurately predict subjects as having abnormal (low) CSF A*β*_1*–*42_ levels indicative of AD risk (0.84 AUC, 0.78 sensitivity, and 0.73 specificity). Refinement of the modeling indicates that only *APOEε4* carrier status and four plasma analytes (CGA, A*β*_1*–*42_, Eotaxin 3, APOE) are required to achieve a high level of accuracy. Furthermore, we show across an independent validation cohort that individuals with predicted abnormal CSF A*β*_1*–*42_ levels transitioned to an AD diagnosis over 120 months significantly faster than those with predicted normal CSF A*β*_1*–*42_levels and that the resulting model also validates reasonably across PET A*β*_1-42_ status (0.78 AUC).

This is the first study to show that a machine learning approach, using plasma protein levels, age and *APOEε4* carrier status, is able to predict CSF A*β*_1*–*42_ status, the earliest risk indicator for AD, with high accuracy.

## 1 Introduction

Alzheimer’s disease (AD) is a terminal neurodegenerative disease that has historically been diagnosed based on “clinically significant” cognitive decline of an individual and exclusion of other conditions. However, it is increasingly recognized that AD is a decades-long neurodegenerative process, with shifts in amyloid *β*^1–42^ (A*β*^1–42^) providing the first indicators of disease development, long before “Alzheimer’s dementia” (significant cognitive decline) is clinically apparent^1–5^.

There is currently no cure or disease-modifying therapy for this terminal illness despite hundreds of clinical trials being conducted since 2002^6,7^. It is hypothesized that the high failure rate of AD trials is in part due to the trials targeting AD patients with significant cognitive impairment, who are therefore in the late stages of the disease and likely have suffered a level of brain tissue loss that cannot be compensated for^8^. Compounding this is the discovery that many patients enrolled in clinical trials were retrospectively found to have normal levels of amyloid and hence did not have AD^9^, with this number as high as 20%^10^. Given these findings, there is a great interest in amyloid screening for clinical trial enrichment in order to recruit individuals at the earliest stages of AD, where intervention is thought to have the greatest chance of success^8^. It would also ensure that included individuals are amyloid positive (i.e. have abnormal levels of amyloid), a necessary precondition for the development of AD. This sort of selective screening is an important precursor for the longer-term goal of population screening for AD^11^.

There are currently two established methods to measure an individual’s amyloid burden: either in vivo in the form of reduced levels of A*β*^1–42^ in the cerebrospinal fluid (CSF) or increased uptake of radioactive tracers that bind selectively to the A*β* fibrillary aggregates by PET imaging. Unfortunately, existing methodologies for measuring an individual’s amyloid levels suffer drawbacks that limit their utility for screening. Lumbar punctures are highly invasive, with this factor alone limiting the applicability of CSF biomarkers for screening. While PET scans are less invasive, they are far more expensive and access to PET scanning facilities is limited in some regions. Despite this, many current trials that target amyloid now require positive amyloid imaging at baseline to ensure accurate diagnosis, a cost-intensive process^7^.

Despite their invasiveness, recent studies have found evidence that changes in CSF A*β*^1–42^ may indicate AD risk long before these same changes are reflected in PET A*β* imaging^12–14^. Palmqvist et al^13^ have shown compelling evidence that changes in CSF A*β*^1–42^ occur up to a decade before the same signal is found by PET A*β* imaging. These results indicate that CSF may be a more suitable measure for early detection, whereas A*β* PET contributes independent information that is more related to disease progression and downstream pathology.

To bypass the invasiveness of CSF collection, there is a strong interest in finding blood-based markers that yield the same information about amyloid status as would be obtained from CSF. There have been a number of studies which have shown that a blood protein signature can be found that reflects AD brain pathology as measured by PET^15–24^. Of particular interest is the recent study by Nakamura et al. (2018) ^25^, whereby levels of A*β*_1*–*40_, A*β*_1*–*42_ and APP_669*–*771_ in plasma, measured using specialised immunoprecipitation (IP) coupled with Matrix Assisted Laser Desorption/Ionization (MALDI) time-of-flight (TOF) mass spectrometry (MS) (henceforth referred to as IP-MALDI-TOF-MS), were shown in combination to have strong performance (> 0.94 area under the receiver operating characteristic curve (AUC)) in predicting PET A*β*_1*–*42_ status across two cohorts. This combination of biomarkers was also found to be predictive of CSF A*β*^1–42^ status, with an AUC of 0.88 on a smaller subset of patients (n=46). The novel IP-MALDI-TOF-MS method employed by Nakamura et al.^25^ is still in its infancy and it is unclear how easily this will be translated into a clinical setting. Thus, there is still strong interest in finding blood markers for CSF A*β*_1*–*42_ using alternative approaches that rely on more established assays.

Here, we evaluated the ability of proteomic and metabolomic data to predict the levels of CSF A*β*_1*–*42_ using a Random Forest (RF) approach and explore which types of measurements lead to the strongest predictive performance. We then determine the minimal set of features required to achieve comparable predictive performance. Finally, we evaluate the robustness and utility of these predictive models across a held-out validation cohort of individuals with mild cognitive impairment (MCI), demonstrating that subjects with predicted abnormal CSF A*β*_1*–*42_ levels showed a faster rate of cognitive decline (measured by the transition to a clinical AD diagnosis) than those with predicted normal CSF A*β*_1*–*42_ levels.

## 2 Methods

### 2.1 Overview of cohort and measurements

The Alzheimer’s Disease Neuroimaging Initiative (ADNI) is a large, multicenter, longitudinal neuroimaging study, launched in 2004 by the National Institute on Aging, the National Institute of Biomedical Imaging and Bioengineering, the Food and Drug Administration, private pharmaceutical companies, and non-profit organizations.

ADNI is a longitudinal study of older adults, designed to test whether serial magnetic resonance imaging (MRI), positron emission tomography (PET), other biological markers, and clinical and neuropsychological assessment can be combined to measure the progression of MCI and early AD.

The ADNI study protocols were approved by the institutional review boards of all participating sites (http://www.adni-info.org/) and written informed consent was obtained from all participants or authorized representatives. All the analytical methods were performed on the de-identified data and were carried out in accordance with the approved guidelines. Study inclusion criteria and definitions for each diagnosis class have been previously reported in detail^26^. Briefly, individuals diagnosed as AD had to meet the National Institute of Neurological and Communicative Disorders and Stroke–Alzheimer’s Disease and Related Disorders Association criteria for probable AD (McKhann et al. 1984). These individuals had issues with global cognition and memory function and they, or their caretakers, reported significant concerns about their memory. In contrast, individuals with MCI exhibited subjective memory loss (CDR of 0.5 and were at least one standard deviation(SD) below the normal mean of the delayed recall of the Wechsler Memory Scale Logical Memory II) but showed preserved activities of daily living, the absence of dementia and scored 24-30 on the MMSE.

### 2.2 Data preparation

We examined 566 individuals in the ADNI cohort who had baseline measures of age, *APOEε*4 carrier status, 193 protein levels (including homocysteine, *Aβ*_1*–*40_, and *Aβ*_1*–*42_) and a further 190 proteins measured on a Rules-Based Medicine (RBM) platform) and 186 LC-MS/MS metabolites and lipids. After applying previously documented quality control procedures (Supplementary Methods) and removing analytes with more than 15% missingness, 149 proteins and 138 metabolites remained. No samples were removed from the analysis as missingness levels were less than 5%. Any remaining missing data points were imputed using an unsupervised RF approach^27^, with the resulting 289 analytes listed in Supplementary Table 5, showing means, SD, and association across different CSF A*β*^1–42^ status. After quality control, a total of 566 individuals, each with measures for 289 analytes, age, and APOE*ε*4 carrier status, were present in the ADNI cohort (Table 1).

**Table 1.** Demographic characteristics of the ADNI data set separated into training and validation cohorts, corresponding to individuals with and without CSF measures respectively. Columns in each cohort provide a further breakdown into individuals that are cognitively normal (CN), have mild cognitive impairment (MCI) or Alzheimer’s disease (AD). The units of each cell are shown in parentheses in the row names and commonly include number of patients (n) or mean of a given quantity (mean). If a secondary measure (percent (%) or standard deviation (SD)) is also present, it is listed in brackets next to the primary measure.

| Dataset | Training |  |  | Validation |  |  |
| --- | --- | --- | --- | --- | --- | --- |
| Diagnosis | CN | MCI | AD | CN | MCI | AD |
| Number of participants (n) | 58 | 198 | 102 | 0 | 198 | 10 |
| Age (mean, (SD)) | 75.11 (5.77) | 74.35 (7.48) | 74.90 (7.91) | - | 75.13 (7.32) | 73.73 (10.04) |
| Gender; female (n, (%)) | 28 (48.28) | 65 (32.82) | 43 (42.16) | - | 75 (37.88) | 4 (40.00) |
| Years of education (mean, (SD)) | 15.67 (2.78) | 15.81 (2.99) | 15.16 (3.29) | - | 15.48 (3.09) | 14.70 (2.06) |
| *APOE*ε4 carriers (n, (%)) | 5 (8.62) | 106 (53.54) | 71 (69.61) | - | 105 (53.03) | 5 (50.00) |
| CSF $A\beta_{1-42}$ abnormal (n, (%)) | 0 (0) | 146 (73.7) | 94 (92.16) | - | - | - |
| PET imaging (n, (%)) | 28 (48.28) | 71 (35.86) | 9 (8.82) | - | 108 (65.55) | - |

### 2.3 Training and Validation cohorts

The 566 individuals in this study ranged from 54.4-89.6 years of age and could be categorized at baseline by their AD clinical diagnosis as cognitively normal (CN; n = 58), amnestic MCI (n = 396) or probable AD (n = 112). A breakdown of the demographics of the 566 individuals by baseline diagnosis and CSF availability is shown in Table 1.

This cohort was split into training and validation cohorts with 356 and 210 individuals with and without measures of CSF A*β*_1*–*42_, respectively. CSF A*β*_1*–*42_ was measured using the Luminex xMAP platform. The training set was used to build predictive models and evaluate their performance directly using the measured A*β*_1*–*42_ levels while the validation cohort was used to evaluate the generalizability and utility of the model’s predictions. For each cohort, we also considered a subset of individuals for whom A*β*^1–42^ status from PET was available at least one-time point (not just at baseline), either using [^11^]C-Pittsburgh compound B (PiB) and [^18^]F-AV-45 (florbetapir, AV45) tracers, for further validation of our modeling. Further demographic information for these cohorts can be found in Supplementary Tables 1, 2 and 3.

### 2.4 Binary and regression modeling tasks

The primary aim of this work was to produce a model that predicts if an individual’s CSF A*β*_1*–*42_ levels are below the recognized clinical threshold of 192pg/ml for the Luminex platform, indicating an abnormal CSF A*β*_1*–*42_ level, and hence increased AD risk. Given the continuous CSF A*β*_1*–*42_ measures in the ADNI cohort, two approaches were considered

- a ‘regression’ task: learning the continuous CSF A*β*_1*–*42_ levels and thresholding these post-prediction
- a ‘binary’ task: learning the dichotomized CSF A*β*_1*–*42_ status based on clinical thresholds directly.

While both tasks result in a binary classifier, they face different trade-offs. The regression task makes use of the full information in the CSF levels but needs to learn a suitable threshold to convert its continuous predictions into suitable binary labels whereas the binary task only learns from the dichotomized CSF levels. Given these trade-offs, we have investigated both modeling approaches throughout this work.

### 2.5 Statistical modeling

We made use of Random Forests (RF) as the modeling approach to predict CSF A*β*_1*–*42_ levels for both the binary and regression tasks. RFs are a widely-used machine-learning ensemble method that have a number of advantages for the small sample size and disparate types of features observed in the ADNI dataset. RFs are invariant to the scale of the observed features and make few assumptions about the distributions of observed data allowing them to be applied to multiple data modalities easily. It can also detect non-additive relationships between variables without needing them to be included explicitly^28^.

All analysis in this work made use of the RF implementation in the R package *ranger*^29^. *Each forest contained 2000 individual trees, each making use of a random selection of p*^3*/*4^ features, where *p* was the total number of variables used in a given model. These parameter choices were based on recommendations provided in Ishwaran *et al.* (2011)^30^. All other parameters in the *ranger* implementation were set to their default values.

To get an estimate of the performance of our models, we have made use of a nested cross-validation (CV) framework, whereby an inner CV was used to determine model parameters, and the outer CV was used to gain an estimate of the model’s performance on unseen data^31^. In this study, we used 3 repetitions of 3 fold CV for the inner loop and 10 repetitions of 10 fold CV for the outer loop.

As the RF used pre-determined parameter values, only a single parameter had to be determined; the threshold on the continuous regression predictions necessary to generate binary labels. This threshold was selected based on performance in the inner CV loop, using the R package *OptimalCutoffs*^32^ to evaluate six potential cutoff metrics (Supplementary Methods) and selecting the method which maximized the accuracy over all of the test folds from the inner cross validation loops. The best performing cutoff criterion was then used in the current iteration of the outer cross-validation loop and the accuracy, sensitivity, and specificity derived from this threshold was recorded for that fold. While this approach means that a different method could be used to derive the regression threshold for each fold in the outer CV loop, the resulting estimate of performance is unbiased and hence is likely to be more representative of performance on unseen data compared with selecting a threshold based on the entire set of training data.

### 2.6 Measures of model performance

Model performance was summarized by the mean and standard deviation of the area under the Receiver Operating Characteristic (ROC) curve (AUC), accuracy, sensitivity, and specificity from the testing performance across the different cross-validation runs. R^2^ values were also calculated for the regression task. Increases in AUC between models were tested for significance using a one-tailed Wilcoxon signed-rank test. Receiver operating curves were constructed by aggregating all of the test predictions from the outer cross-validation.

### 2.7 Evaluating the importance of different input modalities

The input variables were separated into three classes: a commonly used baseline model (B) including age and *APOEε*4 carrier status; Proteomics (P), which included the 146 analytes measured on the RBM panel as well as homocysteine and plasma A*β*_1*–*40_ and A*β*_1*–*42_; Metabolomics (M), including 138 metabolites and lipids.

Four separate random forests were created using different subsets of these features to determine which were most useful for modeling CSF A*β*_1*–*42_. We denoted these models by the combination of features they included; for example ‘BPM’ refers to a model built using all three classes of features. The best performing model was selected for all subsequent analysis.

### 2.8 Discovery of the smallest set of markers needed for strong predictive performance

After evaluating the impact of the different input modalities, we determine the minimum set of individual analytes necessary to achieve high predictive performance. This was done by treating the number of included features as a parameter to be determined in our nested CV framework. Within each fold of the inner CV loop, we used a recursive feature elimination approach, ranking features according to their *Variable Importance*, the difference in the prediction error on the out-of-bag data when a given feature was permuted and unpermuted^28^ and removed the lowest ranking features in a stepwise fashion. The AUC of the resulting RF was recorded, and the procedure was repeated over increasingly smaller subsets of features until no features were left to be removed. After the inner CV loop finishes, we determine the number of features that achieved the optimal trade-off between model complexity and performance by selecting the smallest subset of features that achieved within 4% of the maximal observed AUC. A model using this subset of features was then trained on all training folds of the outer CV loop and evaluated on the test fold. Again, by determining the number of features to include within our nested cross-validation framework, we are able to determine an unbiased estimate of the model’s expected performance over unseen data.

### 2.9 Survival Analysis

Survival analysis was conducted to determine if the rate of conversion from MCI to AD was different between those with predicted low and normal CSF A*β*_1*–*42_ levels, enabling us to determine if our predictions lead to useful clinical outcomes in the validation cohort.

Four separate analyses were performed, using the:

1. measured CSF status on the training set (n=198)
2. predicted CSF status from B model, the standard baseline, in the validation set (n=198)
3. predicted CSF status from BP model, the best performing model, in the validation set (n=198)
4. predicted CSF status from BP _*fs*_ model, the most parsimonious model, in the validation set (n=198),

where BP _*fs*_ is the feature selected model with the smallest set of features. For each analysis, we have examined the hazard ratios using Cox regression and used log-rank tests to compare the survival distribution of low/normal CSF A*β*_1*–*42_ stratifications in the four analyses, as well to compare equivalence between the actual and predicted stratifications.

### 2.10 Validation performance over PET A*β*_1*–*42_ status

In order to further validate our model, we have examined the ability of our model to differentiate PET A*β*_1*–*42_ abnormal and normal status. While it is known that A*β*_1*–*42_ status from PET can differ from that observed in CSF, measurements from the two modalities are correlated and should be very similar for individuals who are not close to the cutoff indicating pathology. This provides analysis provides further evidence of our model’s ability to determine A*β*_1*–*42_ status in individuals in the validation cohort, where CSF measurements are not available.

Given that only a limited number of individuals had associated measures of PET imaging at baseline (n=18 and 27 for training and validation cohorts respectively), we have made use of the earliest PET image available, leaving us with 108 and 68 individuals in the training and validation cohort to evaluate. The threshold for abnormality was defined as an SUVR of 1.5 and 1.11 for PET images using PiB and AV45 tracers respectively. The mean number of years past baseline that a scan was taken was 3.07 and 2.97 years for training and validation cohorts respectively.

The use of imaging at non-baseline times assumes that differences between the baseline and time that the image was taken are relatively small (which may be reasonable assuming a slow rate of A*β*_1*–*42_ accumulation) and that few individuals are close to the defined threshold for abnormality. If these assumptions do not hold, it is likely to worsen predictive performance, making this analysis somewhat conservative.

## 3. Results

### 3.1 Models utilizing protein levels accurately predict CSF positivity

We evaluated the ability of blood-based biomarkers to predict CSF *Aβ*_1*–*42_ normal/abnormal status using RFs trained using different subsets of input variables, treating the modeling of CSF A*β*_1*–*42_ as either a regression or binary task. Summaries of the performance metrics from the resulting models are shown in Table 2 with their corresponding ROC curves shown in Figure 1.

**Figure 1.**
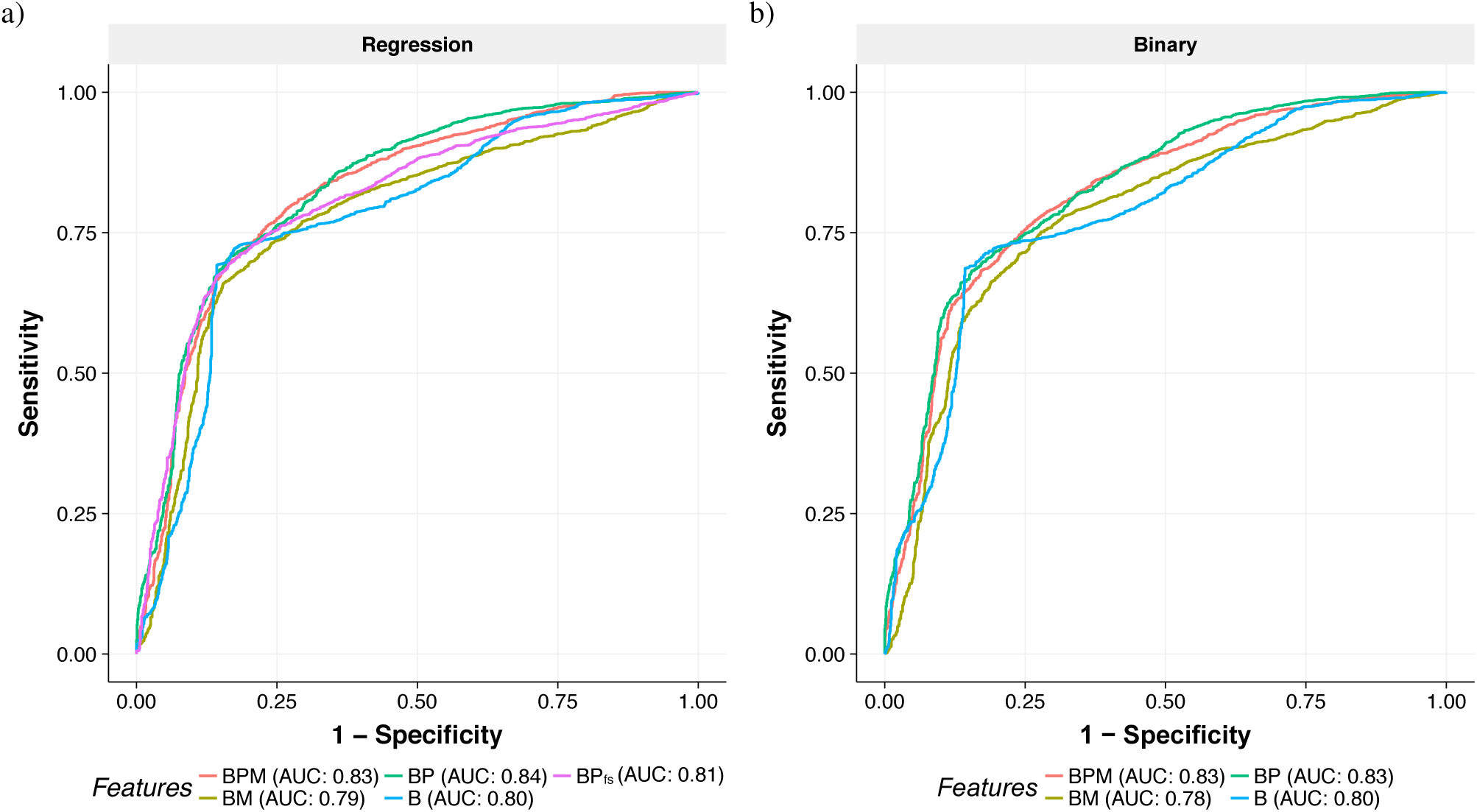
ROC curves comparing different sets of features to determine predictive value for the a) regression, and b) binary tasks. Different colours of lines correspond to different feature sets, (B) baseline model (age and *APOEε*4 carrier status), (P) Proteomics, (M) Metabolomics, with corresponding AUCs indicating in the legend under each plot. The model BP _*fs*_ in subplot a) indicates the performance of the RF using feature selection.

**Table 2.** Mean and standard deviation (in parentheses) of performance metrics (area under the receiver operator curve, AUC; accuracy, Acc; sensitivity, Sens; specificity, Spec and *R*^2^ for the regression models) for the different Random Forest models using different feature sets across all cross-validation folds. Left and right halves are for the Regression and Binary tasks respectively. Bold faced text on AUCs indicates the best performing model or those that are statistically equivalent (via a Wilcox rank signed test, with a Bonferroni-corrected significance threshold of 0.05/5=0.01). Features sets describe combinations of (B) baseline model (age and *APOEε*4 carrier status), (P) Proteomics, (M) Metabolomics.

| Feature set | Regression |  |  |  |  | Binary |  |  |  |
| --- | --- | --- | --- | --- | --- | --- | --- | --- | --- |
| | AUC | $R^2$ | Acc | Sens | Spec | AUC | Acc | Sens | Spec |
| BPM | 0.830 (0.08) | 0.274 (0.11) | **0.771 (0.07)** | 0.784 (0.09) | **0.748 (0.13)** | **0.826 (0.07)** | **0.766 (0.06)** | **0.884 (0.06)** | 0.536 (0.14) |
| BM | 0.788 (0.08) | 0.209 (0.13) | 0.744 (0.08) | 0.779 (0.09) | 0.679 (0.16) | 0.784 (0.08) | 0.737 (0.08) | 0.855 (0.08) | 0.507 (0.15) |
| BP | **0.839 (0.07)** | **0.293 (0.11)** | **0.765 (0.08)** | 0.782 (0.10) | **0.731 (0.13)** | **0.834 (0.07)** | **0.769 (0.07)** | **0.882 (0.08)** | 0.552 (0.14) |
| B | 0.795 (0.08) | **0.275 (0.14)** | 0.746 (0.06) | 0.751 (0.08) | **0.747 (0.15)** | 0.800 (0.07) | 0.745 (0.07) | 0.731 (0.09) | **0.780 (0.11)** |
| BP<sub>fs</sub> | 0.812 (0.08) | 0.270 (0.14) | 0.749 (0.07) | **0.807 (0.10)** | 0.638 (0.15) |  |  |  |  |

We observed strong overall predictive performance for both the regression and binary tasks within our cross-validation framework. All sets of features outperformed the base model of age and *APOEε*4 carrier status with BP based models leading to the highest AUC of 0.84 and 0.83 for the regression and binary tasks respectively. The standard deviation for the AUC was relatively high (7-8%), likely due to the noise inherent in both the analytes being used for prediction as well as in the CSF A*β*_1*–*42_ measurements.

The BP models resulted in a mean *R*^2^ of 0.29 for the regression task. The automatically derived threshold for this regression RF yielded a mean accuracy of 0.77, with a sensitivity of 0.78 and a specificity of 0.73. Across the 100 cross-validation runs, the chosen threshold ranged from 164pg/ml to 194pg/ml with a median of 185pg/ml.

Similar AUC and accuracy could be observed for learning the dichotomized CSF labels directly (e.g 0.83 AUC, 0.77 accuracy for the BP model). For the binary task, a slight drop in both AUC, as well as an altered trade-off between sensitivity and specificity, was observed across all different feature sets compared to the regression task. Given this, we chose to focus on the regression model for much of the follow-up analysis.

While all models making use of blood analytes outperformed the base model of age and APOE*ε*4, models that made use of the protein level measurements consistently achieved the strongest predictive performance, whereas metabolites appeared to be of limited utility. In both the regression and binary tasks, models containing metabolites and proteomic data (BPM) achieved equivalent or worse AUCs than models containing only the proteomic data (BP). Furthermore, we observed that the use of the base features and metabolites alone (BM) lead to decreased performance compared to the baseline model, indicating that the set of measured metabolites may have contributed little predictive information or may have been too noisy to be useful for predicting CSF status. These findings are in contrast to the previously reported utility of metabolites in predicting PET A*β*^1–42^ positivity^22^.

While the results presented in this section include clinically diagnosed AD individuals, who are almost all CSF A*β*_1*–*42_ positive, it is worth considering only ‘pre-clinical’ individuals as this may be more relevant for selective screening in drug trials. Evaluating our model’s performance on CN and MCI individuals only, we find that similarly strong predictive performance can be obtained (Supplementary Table 4, Supplementary Figure 1, 0.80 AUC, 0.77 accuracy for the BP model) supporting our primary findings that plasma protein levels can be utilized to predict amyloid pathology status.

To ensure that our imputation procedure did not bias our results, we also built similar models using only complete cases after applying more stringent quality control (removal of plasma analytes where more than 1% of measurements were missing), obtaining similar AUCs of 0.81 for the regression and binary tasks (Supplementary Figure 2).

### 3.2 Strong predictive performance is maintained using only four proteins

The models described so far used all (*>* 140) available features in this dataset. In practice, measuring hundreds of analytes is costly, negating a key advantage of using blood biomarkers for screening. Given this, we have applied feature selection to the BP regression model to identify the smallest number of features that still achieved high predictive performance. Within cross-validation, we find that the average performance of this feature selection approach, denoted BP _*fs*_, yields an AUC, sensitivity, and specificity of 0.81, 0.81 and 0.63. The number of features selected in the model ranges from 2 to 15, with a median of 5 features included.

When applying this feature selection procedure to the entire set of training data, we identified a subset of four plasma analytes as well as APOE4 genotype status critical for model performance: Chromogranin-A (CGA), A*β*^1–42^ (AB42), Eotaxin 3, and Apolipoprotein E (APOE). This combination of protein levels, together with *APOEε*4 is denoted as BP5. Figure 2 indicates how each variable influences the model predictions after we have accounted for the influence of the other four variables. As expected, the strongest relationship with CSF A*β*_1*–*42_ is with *APOEε*4 carrier status, where being a carrier (*APOEε*4 = 1) leads to a low predicted A*β*_1*–*42_ level. While the relationships between the proteins and CSF A*β*_1*–*42_ are non-linear (a common outcome given the nature of RFs), the overall correlation with CSF A*β*_1*–*42_ is positive for CGA, Plasma A*β*_1*–*42_, and APOE protein levels and negative for Eotaxin 3.

**Figure 2.**
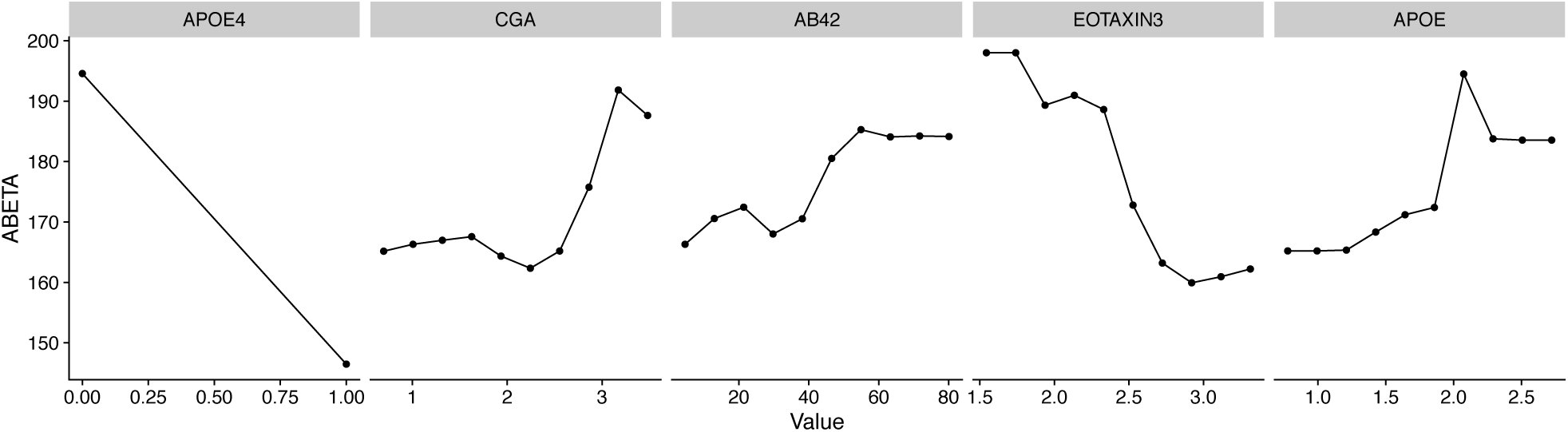
Partial dependency plots of the five features selected from the full BP model using a recursive feature elimination approach. Each subplot shows how the variation of a specific feature impacts that predicted levels of CSF A*β*_1*–*42_ assuming the other four features are fixed.

### 3.3 Validation of clinical utility

To demonstrate the utility of our modeling on unseen data, we conducted a survival analysis over the validation cohort (n=198), evaluating the probability of baseline MCI individuals transitioning to AD diagnosis over 120 months, stratified by predicted CSF *Aβ*_1*–*42_ status from either the B, BP or reduced BP5 model. These survival distributions could then be compared to those of the real A*β*_1*–*42_ status observed over the training cohort. Given the demographic similarity of the two cohorts, we would expect to see strong similarities in rates of conversion.

From Figure 3, we observed that in all cases, the predicted low CSF A*β*_1*–*42_ group transitioned to AD significantly faster than the A*β*_1*–*42_ normal group. Comparing the predictions from the BP, BP5 and B models on the validation cohort to the actual CSF A*β*_1*–*42_ status on the training cohort, we find that there is no significant difference between the survival distributions for either the normal (log-rank test *p* = 0.19, 0.2, 0.21) or abnormal (log-rank test *p* = 0.97, 0.31, 0.23) survival distributions, respectively, reflecting the overlapping confidence intervals of the hazard ratios. However, it can be observed that due to differences in the thresholding of the A*β*_1*–*42_ levels, fewer individuals are deemed as CSF A*β*_1*–*42_ ‘normal’ in the actual data (n=53), compared with any of the three models applied to the validation datasets (n=95, 73, and 71 for BP, BP5, and B models respectively), highlighting the well-recognized issues of defining standardized cutoff values across studies^33^. The significant differences in conversion rates between the predicted normal/low strata, especially from the more parsimonious BP5 model, together with their similarity to the survival distributions of the actual CSF measures, provide strong evidence that our blood-based model can help stratify individuals based on their risk of developing clinical AD (Table 1).

**Figure 3.**
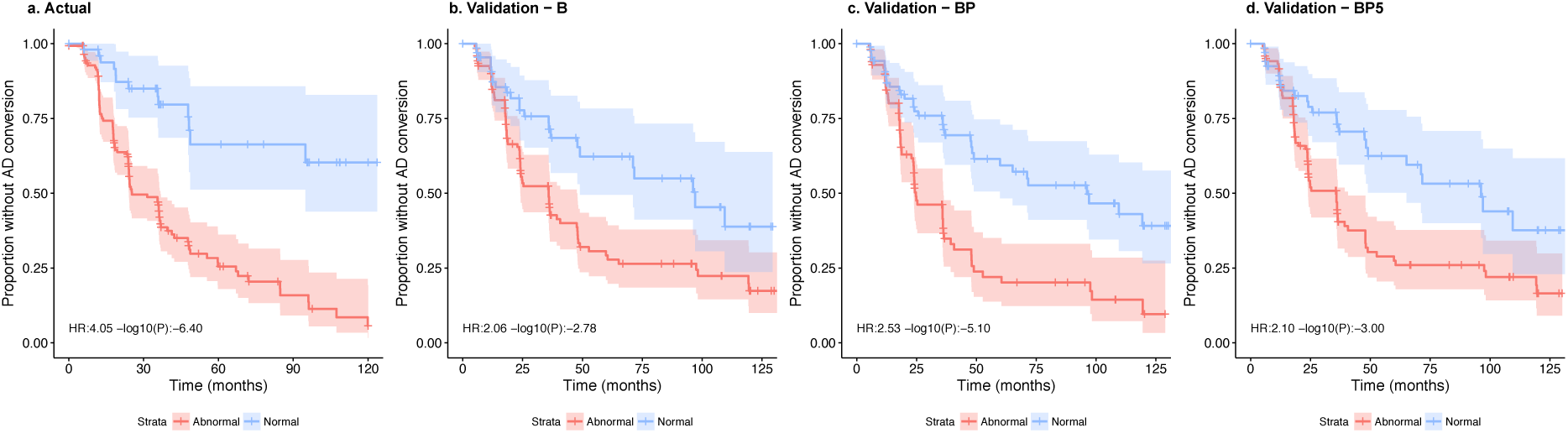
Kaplan-Meier curves for a) the training cohort stratified by actual CSF A*β*_1*–*42_ status, and the validation cohort stratified by predicted CSF A*β*_1*–*42_ status from the b) B model, c) BP model and d) BP5 model. The bands along the curves represent the 95% confidence intervals. Hazards ratios and 95% confidence intervals for the abnormal group compared to the normal are shown in the bottom left of each subplot. In all cases, the low CSF A*β*^1–42^ group transitioned to AD diagnosis significantly faster then the normal group (*p* = 3.97 ×10^*–*7^, 7.89× 10^*–*6^, 9.96 ×10^*–*4^, 1.65× 10^*–*3^ for the four plots left-right). CN individuals were not included in this analysis because there were no CN individuals present in the validation cohort.

### 3.4 Concordance with PET A*β*_1*–*42_ status

To further validate and quantify our model’s performance, we have explored the relationship between the predicted CSF A*β*_1*–*42_ scores and PET imaging status. Confirming that the PET and CSF A*β*_1*–*42_ status are correlated, we find that they differed in only 7 out of 108 individuals for whom both CSF and PET amyloid status were available. As such, evaluating our model against the PET A*β*_1*–*42_ status should provide a conservative estimate for the AUC on the validation cohort, despite the lack of CSF measures.

The resulting ROC curves in Figure 4 provide further evidence that the BP and BP5 models are able to predict A*β*_1*–*42_ status, with AUCs against PET A*β*_1*–*42_ on the validation cohort of 0.78 and 0.8 for the BP and BP5 models respectively. These results are similar to those from predicting CSF status from the training data (Figure 1), with a small expected drop due to the inherent differences between CSF and PET amyloid. Interestingly, we observe stronger performance for the reduced BP5 model compared to the full BP model, with both models significantly improving upon the baseline model of age and *APOEε*4 status.

**Figure 4.**
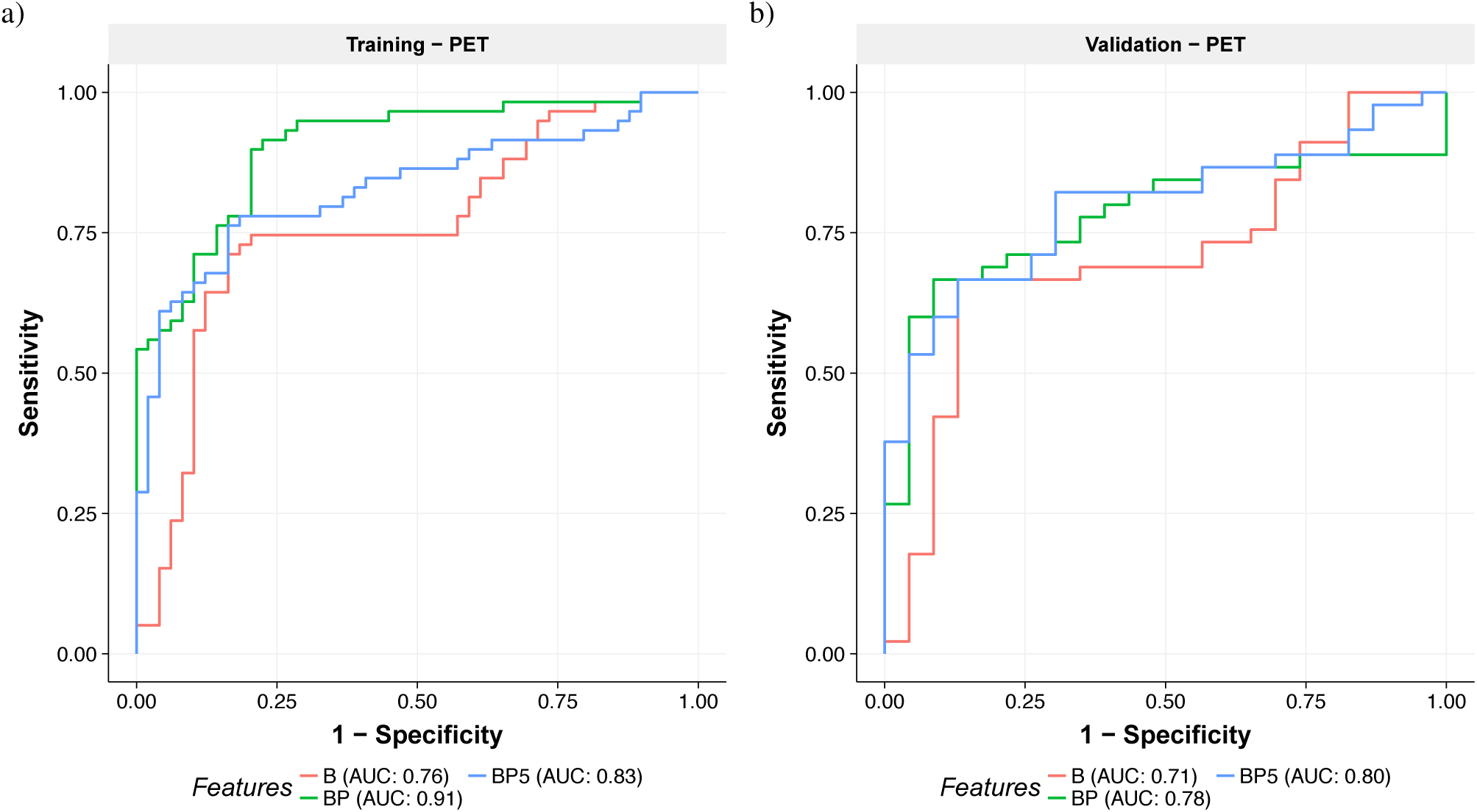
ROC curves comparing different sets of features to determine PET-based A*β*_1*–*42_ status on the a) training and b) validation cohorts. Different colours of lines correspond to different feature sets with corresponding AUCs indicating in the legend under each plot. Results in the training cohort are more useful as a measure of similarity between the tasks of predicting CSF and PET A*β*_1*–*42_ status given that this was the data used to train the CSF model and hence the AUC are upwardly biased, especially for more complex models (e.g. BP).

## 4 Discussion

The most positive results from AD trials to date have been found in patients with early forms of the disease, leading to an increasing awareness that treatments are likely to be most successful if applied at the earliest stages of AD^8^. Some AD clinical trials are enriching pre-symptomatic AD individuals with PET screening. However, recent findings that shifts in CSF amyloid can be observed up to a decade before those from PET may indicate that CSF positive individuals are even more suitable for clinical trial enrichment ^34^. Direct measurement of CSF biomarkers is too invasive to be used in such a screening test^35^ motivating the development of a minimally-invasive, low-cost solution that provides the same type of information to overcome these limitations.

This current study evaluates the utility of a blood-based signature of CSF A*β*_1*–*42_ status using a Random Forest approach. We demonstrated that CSF A*β*^1–42^ normal/abnormal status using age, *APOEε4* carrier status, and protein levels can be predicted with a high AUC, sensitivity and specificity of 0.84, 0.78 and 0.73 respectively. Compared to the base model (age and *APOEε*4 genotype) the inclusion of the plasma analytes improved the performance (AUC) by 6%. To make the model more suitable for clinical application, we identified four plasma analytes which, together with *APOEε4* carrier status, still achieved a high AUC, sensitivity, and specificity of 0.81, 0.81 and 0.64 respectively. These predictive models were then validated on a separate cohort of individuals to demonstrate that MCI subjects with predicted abnormal CSF A*β*_1*–*42_ (low) levels transitioned to an AD diagnosis at a significantly higher rate than those predicted with normal CSF A*β*_1*–*42_ levels. Furthermore, these rates were similar to those observed in a demographically similar cohort of MCIs using actual CSF A*β*_1*–*42_ levels. This is a strong validation of our modeling as the blood-based biomarkers for CSF A*β*_1*–*42_ status is only useful if they can replicate the behavior of the actual A*β*_1*–*42_ status for clinically relevant endpoints for individuals that were not used to build the predictive model. Strong predictive power of PET A*β*_1*–*42_ status on the validation cohort provides further evidence for the generalizability and robustness of our modeling.

A number of studies have previously investigated the use of blood analytes to predict the burden of amyloid in the neocortex, as measured by PET ^15, 16, 18–20, 22, 23^. Some of these studies showed similar performance metrics to those reported in this work (> 0.80 AUCs^15,23–25^ or > 0.78 accuracy^17^), indicating that prediction of PET and CSF *Aβ*_1*–*42_ status are of similar difficulty. PET A*β* is directly related to brain fibrillar amyloid, whereas CSF amyloid is a marker of soluble A*β*_1*–*42_ and they may, therefore, give different insights into AD progression and mechanisms. For example, CSF A*β*_1*–*42_ has been shown to be associated with *APOEε4* whereas PET A*β*_1*–*42_ has been shown to have a greater association with tau^36^. Thus, the development of a blood-based screening test for CSF A*β*_1*–*42_ levels is a complementary approach to existing blood-based biomarkers of PET amyloid status

Of the above studies, the study by Nakamura et. al.^25^ showed a very high AUC in discovery and validation datasets for PET A*β*_1*–*42_ status (AUC 0.94 and 0.96 respectively) as well as a strong performance for predicting abnormalities in CSF A*β*_1*–*42_ levels (AUC 0.88%), in a small cohort (n=46) of their validation set. While these results are promising, the automation of the novel technique used (IP-MALDI-TOF-MS), and hence transfer to a clinical setting, is non-trivial, motivating the search for complementary approaches. The protein signature presented in this study, based on a multiplex immunoassay, is likely to require a far shorter timeframe for clinical translation given the high level of automation that already exists for multiplex immunoassays, and that biomarkers from such platform have already been used in commercially available diagnostic tests that have been approved by the FDA.

The use of metabolites appeared to be of limited utility for predicting CSF A*β*_1*–*42_. In both the regression and binary tasks, models containing metabolites achieved equivalent or worse AUCs than models without. These findings can be contrast with the utility of metabolites in predicting PET A*β*_1*–*42_ positivity^22^ and their association with AD more broadly^37^. Alternative methods for integrating this source of data^38^ may be required in order to find robust associations with CSF A*β*_1*–*42_ status.

The subset of features used in our BP5 model included *APOEε4* genotype and plasma levels of Chromogranin-A (CGA),Eotaxin 3, A*β*^1–42^ (AB42), and Apolipoprotein E (APOE). Several of these identified proteins have known associations with Alzheimer’s disease. Unsurprisingly, the levels of plasma APOE are associated with CSF amyloid levels. *APOEε4* is the strongest genetic risk factor for AD. APOE is involved in the clearance of [inline]and there is a strong relationship between APOE*ε*4 genotype and APOE plasma levels, where APOE*ε*4 carriers have lower plasma levels^42, 43^. Plasma A*β*^1–42^ showed a positive relationship in our model for CSF A*β*_1*–*42_, in line with a prior observation^44^. This is interesting as the link between alterations of A*β*^1–42^ levels in the blood and the progression of the disease is still controversial and studies assessing the A*β*^1–42^ concentration in blood of AD patients have produced conflicting results^44–50^. Chromogranin A (CGA) is associated with synaptic function and has traditionally been used as an indicator of neuroendocrine tumors^51^. More recent work has shown that CGA has a degree of co-localisation with amyloid plaques in the brain^52, 53^. However, levels of CGA in the CSF and blood serum do not appear to be correlated^54^ and serum CGA has not previously been linked to AD. Eotaxin 3, also known as C-C chemokine ligand 26 (CCL26), plays an important role in the innate immune system and has been found to be dysregulated in AD patients^55^. CSF Eotaxin 3 has been shown to be significantly elevated in patients with prodromal AD, however, Eotaxin 3 levels in plasma or the CSF has not been shown to correlate with rates of disease progression^55, 56^

This study has several limitations. The training and validation cohorts are both composed of individuals in the ADNI study and thus all measures were conducted on the same platforms. Hence further cross-cohort and cross-platform replication is required. This remains an ongoing issue within the development of all AD biomarkers relating to early screening and requires significant future investment^57^. Furthermore, the current cohort is neuropathology biased, i.e. 84% of the cohort have MCI or AD, and thus likely to have neuronal damage, potentially confounding the analysis of CSF A*β*_1*–*42_ status. Finally, it needs to be noted that there are other medical conditions that are known to affect CSF A*β*_1*–*42_ levels and it is unclear whether these affect any of the patients in our cohort.

The early identification of AD disease is paramount and a major global focus as the success of disease-modifying or preventative therapies in AD may depend on detecting the earliest signs of abnormal amyloid-beta load. The differences between CSF A*β*_1*–*42_ and PET A*β*_1*–*42_ in preclinical stages of AD are likely to have implications for clinical trial enrichment. Blood-based biomarkers of amyloid can serve as the first step in a multistage screening procedure, similar to those that have been clinically-implemented in cancer, cardiovascular disease, and infectious diseases^57^. In-conjunction with biomarkers for neocortical amyloid burden, the CSF A*β*_1*–*42_ biomarkers presented in this work may help yield a cheap, non-invasive tool for both improving clinical trials targeting amyloid and population screening.

## 5 Acknowledgments

Data collection and sharing for this project was funded by the Alzheimer’s Disease Neuroimaging Initiative (ADNI) (National Institutes of Health Grant U01 AG024904) and DOD ADNI (Department of Defense award number W81XWH-12-2-0012). ADNI is funded by the National Institute on Aging, the National Institute of Biomedical Imaging and Bioengineering, and through generous contributions from the following: AbbVie, Alzheimer’s Association; Alzheimer’s Drug Discovery Foundation; Araclon Biotech; BioClinica, Inc.; Biogen; Bristol-Myers Squibb Company; CereSpir, Inc.; Cogstate; Eisai Inc.; Elan Pharmaceuticals, Inc.; Eli Lilly and Company; EuroImmun; F. Hoffmann-La Roche Ltd and its affiliated company Genentech, Inc.; Fujirebio; GE Healthcare; IXICO Ltd.; Janssen Alzheimer Immunotherapy Research & Development, LLC.; Johnson & Johnson Pharmaceutical Research & Development LLC.; Lumosity; Lundbeck; Merck & Co., Inc.; Meso Scale Diagnostics, LLC.; NeuroRx Research; Neurotrack Technologies; Novartis Pharmaceuticals Corporation; Pfizer Inc.; Piramal Imaging; Servier; Takeda Pharmaceutical Company; and Transition Therapeutics. The Canadian Institutes of Health Research is providing funds to support ADNI clinical sites in Canada. Private sector contributions are facilitated by the Foundation for the National Institutes of Health (www.fnih.org). The grantee organization is the Northern California Institute for Research and Education, and the study is coordinated by the Alzheimer’s Therapeutic Research Institute at the University of Southern California. ADNI data are disseminated by the Laboratory for Neuro Imaging at the University of Southern California. This work was supported by IBM. We would like to thank Dr. Matthew Downton, Dr. Annalisa Swan and Dr. Anna Trigos for helpful feedback on the manuscript.

## 7 Author Contributions Statement

B.G, B.F, and N.F. designed the study, B.G., C.S. and B.F. analyzed the data and ran all experiments, B.G. made the figures, B.G., C.S. and N.F. wrote the manuscript. All authors interpreted the data and critically revised the manuscript.

## 8 Consortium Members

### 8.1 Alzheimer’s Disease Neuroimaging Initiative - ADNI

Michael W. Weiner^8^, Paul Aisen^9^, Ronald Petersen^10^, Clifford R. Jack, Jr.^11^, William Jagust^12^, John Q. Trojanowki^13^, Arthur W. Toga^14^, Laurel Beckett^15^, Robert C. Green^16^, Andrew J. Saykin^17^, John Morris^18^, Leslie M. Shaw^13^, Jeffrey Kaye^19^, Joseph Quinn^20^, Lisa Silbert^20^, Betty Lind^20^, Raina Carter^19^, Sara Dolen^19^, Lon S. Schneider^19^, Sonia Pawluczyk^19^, Mauricio Beccera^19^, Liberty Teodoro^14^, Bryan M. Spann^14^, James Brewer^20^, Helen Vanderswag20, Adam Fleisher20, Judith Heidebrink^21^, Joanne L. Lord^21^, Sara S. Mason^11^, Colleen S. Albers^11^, David Knopman^11^, Kris Johnson^11^, Rachelle S. Doody^22^, Javier Villanueva-Meyer^22^, Munir Chowdhury^22^, Susan Rountree^22^, Mimi Dang^22^, Yaakov Stern^23^, Lawrence S. Honig^23^, Karen L. Bell^23^, Beau Ances^23^, John C. Morris^23^, Maria Carroll^23^, Mary L. Creech^23^, Erin Franklin^23^, Mark A. Mintun^18^, Stacy Schneider^18^, Angela Oliver^18^, Daniel Marson^24^, Randall Griffth^24^, David Clark^24^, David Geldmacher^24^, John Brockington^24^, Erik Roberson^24^, Marissa Natelson Love^24^, Hillel Grossman^25^, Effie Mitsis^25^, Raj C. Shah^26^, Leyla deToledo-Morrell^26^, Ranjan Duara^27^, Daniel Varon^27^, Maria T. Greig^27^, Peggy Roberts^27^, Marilyn Albert^28^, Chiadi Onyike^28^, Daniel D’Agostino^28^, Stephanie Kielb^28^, James E. Galvin^29^, Brittany Cerbone^29^, Christina A. Michel^29^, Dana M. Pogorelec^29^, Henry Rusinek^29^, Mony J de Leon^29^, Lidia Glodzik^29^, Susan De Santi^29^, P. Murali Doraiswamy^30^, Jeffrey R. Petrella^30^, Salvador Borges-Neto^30^, Terence Z. Wong^30^, Edward Coleman^30^, Charles D. Smith^31^, Greg Jicha^31^, Peter Hardy^31^, Partha Sinha^31^, Elizabeth Oates^31^, Gary Conrad^31^, Anton P. Porsteinsson^32^, Bonnie S. Goldstein^32^, Kim Martin^32^, Kelly M. Makino^32^, Saleem Ismail^32^, Connie Brand^32^, Ruth A. Mulnard^33^, Gaby Thai^33^, Catherine Mc-Adams-Ortiz^33^, Kyle Womack^34^, Dana Mathews^34^, Mary Quiceno^34^, Allan I. Levey^35^, James J. Lah^35^, Janet S. Cellar^35^, Jeffrey M. Burns^36^, Russell H. Swerdlow^36^, William M. Brooks^36^, Liana Apostolova^37^, Kathleen Tingus^37^, Ellen Woo^37^, Daniel H.S. Silverman^37^, Po H. Lu^37^, George Bartzokis^37^, Neill R Graff-Radford^38^, Francine Parftt^38^, Tracy Kendall^38^, Heather Johnson^38^, Martin R. Farlow^17^, Ann Marie Hake^17^, Brandy R. Matthews^17^, Jared R. Brosch^17^, Scott Herring^17^, Cynthia Hunt^17^, Christopher H. van Dyck^39^, Richard E. Carson^39^, Martha G. MacAvoy^39^, Pradeep Varma^39^, Howard Chertkow^40^, Howard Bergman^40^, Chris Hosein^40^, Sandra Black^41^, Bojana Stefanovic^41^, Curtis Caldwell^41^, Ging-Yuek Robin Hsiung^42^, Howard Feldman^42^, Benita Mudge^42^, Michele Assaly^42^, Elizabeth Finger^43^, Stephen Pasternack^43^, Irina Rachisky^43^, Dick Trost^43^, Andrew Kertesz^43^, Charles Bernick^44^, Donna Munic^44^, Marek-Marsel Mesulam^45^, Kristine Lipowski^45^, Sandra Weintraub^45^, Borna Bonakdarpour^45^, Diana Kerwin^45^, Chuang-Kuo Wu^45^, Nancy Johnson^45^, Carl Sadowsky^46^, Teresa Villena^46^, Raymond Scott Turner^47^, Kathleen Johnson^47^, Brigid Reynolds^47^, Reisa A. Sperling^48^, Keith A. Johnson^48^, Gad Marshall^48^, Jerome Yesavage^49^, Joy L. Taylor^49^, Barton Lane^49^, Allyson Rosen^49^, Jared Tinklenberg^49^, Marwan N. Sabbagh^50^, Christine M. Belden^50^, Sandra A. Jacobson^50^, Sherye A. Sirrel^50^, Neil Kowall^51^, Ronald Killiany^51^, Andrew E. Budson^51^, Alexander Norbash^51^, Patricia Lynn Johnson^51^, Thomas O. Obisesan^52^, Saba Wolday^52^, Joanne Allard^52^, Alan Lerner^53^, Paula Ogrocki^53^, Curtis Tatsuoka^53^, Parianne Fatica^53^, Evan Fletcher^54^, Pauline Maillard^54^, John Olichney^54^, Charles DeCarli^54^, Owen Carmichael^54^, Smita Kittur^55^, Michael Borrie^56^, T-Y Lee^56^, Rob Bartha^56^, Sterling Johnson^57^, Sanjay Asthana^57^, Cynthia M. Carlsson^57^, Steven G. Potkin^57^, Adrian Preda^57^, Dana Nguyen^57^, Pierre Tariot^58^, Anna Burke^58^, Nadira Trncic^58^, Adam Fleisher^59^, Stephanie Reeder^59^, Vernice Bates^60^, Horacio Capote^60^, Michelle Rainka^60^, Douglas W. Scharre^61^, Maria Kataki^61^, Anahita Adeli^61^, Earl A. Zimmerman^62^, Dzintra Celmins^62^, Alice D. Brown^62^, Godfrey D. Pearlson^63^, Karen Blank^63^, Karen Anderson^63^, Laura A. Flashman^64^, Marc Seltzer^64^, Mary L. Hynes^64^, Robert B. Santulli^64^, Kaycee M. Sink^65^, Leslie Gordineer^65^, Je D. Williamson^65^, Pradeep Garg^65^, Franklin Watkins^65^, Brian R. Ott^66^, Henry Querfurth^66^, Geffrey Tremont^66^, Stephen Salloway^67^, Paul Malloy^67^, Stephen Correia^67^, Howard J. Rosen^68^, Bruce L. Miller^68^, David Perry^68^, Jacobo Mintzer^69^, Kenneth Spicer^69^, David Bachman^69^, Nunzio Pomara^70^, Raymundo Hernando^70^, Antero Sarrael^70^, Norman Relkin^71^, Gloria Chaing^71^, Michael Lin^71^, Lisa Ravdin^71^, Amanda Smith^72^, Balebail Ashok Raj^72^, Kristin Fargher^72^

8 Magnetic Resonance Unit at the VA Medical Center and Radiology, Medicine, Psychiatry and Neurology, University of California, San Francisco, USA.

9 San Diego School of Medicine, University of California, California, USA.

10 Mayo Clinic, Minnesota, USA.

11 Mayo Clinic, Rochester, USA.

12 University of California, Berkeley, USA.

13 University of Pennsylvania, Pennsylvania, USA.

14 University of Southern California, California, USA.

15 University of California, Davis, California, USA.

16 MPH Brigham and Women’s Hospital/Harvard Medical School; Massachusetts, USA.

17 Indiana University, Indiana, USA.

18 Washington University St. Louis, Missouri, USA.

19 Oregon Health and Science University, Oregon, USA.

20 University of California–San Diego, California, USA.

21 University of Michigan, Michigan, USA.

22 Baylor College of Medicine, Houston, State of Texas, USA.

23 Columbia University Medical Center, South Carolina, USA.

24 University of Alabama – Birmingham, Alabama, USA.

25 Mount Sinai School of Medicine, New York, USA.

26 Rush University Medical Center, Rush University, Illinois, USA.

27 Wien Center, Florida, USA.

28 Johns Hopkins University, Maryland, USA.

29 New York University, NY, USA.

30 Duke University Medical Center, North Carolina, USA.

31 University of Kentucky, Kentucky, USA.

32 University of Rochester Medical Center, NY, USA.

33 University of California, Irvine, California, USA.

34 University of Texas Southwestern Medical School, Texas, USA.

35 Emory University, Georgia, USA.

36 University of Kansas, Medical Center, Kansas, USA.

37 University of California, Los Angeles, California, USA.

38 Mayo Clinic, Jacksonville, USA.

39 Yale University School of Medicine, Connecticut, USA.

40 VMcGill University, Montreal-Jewish General Hospital, Canada.

41 Sunnybrook Health Sciences, Ontario, USA.

42 U.B.C. Clinic for AD & Related Disorders, Canada.

43 Cognitive Neurology - St. Joseph’s, Ontario, USA.

44 Cleveland Clinic Lou Ruvo Center for Brain Health, Ohio, USA.

45 Northwestern University, USA.

46 Premiere Research Inst (Palm Beach Neurology), USA.

47 Georgetown University Medical Center, Washington D.C, USA.

48 Brigham and Women’s Hospital, Massachusetts, USA.

49 Stanford University, California, USA.

50 Banner Sun Health Research Institute, USA.

51 Boston University, Massachusetts, USA.

52 Howard University, Washington D.C, USA.

53 Case Western Reserve University, Ohio, USA.

54 University of California, Davis – Sacramento, California, USA.

55 Neurological Care of CNY, USA.

56 Parkwood Hospital, Pennsylvania, USA.

57 University of Wisconsin, Wisconsin, USA.

58 University of California, Irvine – BIC, USA.

59 Banner Alzheimer’s Institute, USA.

60 Dent Neurologic Institute, NY, USA.

61 Ohio State University, Ohio, USA.

62 Albany Medical College, NY, USA.

63 Hartford Hospital, Olin Neuropsychiatry Research Center, Connecticut, USA.

64 Dartmouth-Hitchcock Medical Center, New Hampshire, USA.

65 Wake Forest University Health Sciences, North Carolina, USA.

66 Rhode Island Hospital, state of Rhode Island, USA.

67 Butler Hospital, Providence, Rhode Island, USA.

68 University of California, San Francisco, USA.

69 Medical University South Carolina, USA.

70 Nathan Kline Institute, Orangeburg, New York, USA.

71 Cornell University, Ithaca, New York, USA.

72 USF Health Byrd Alzheimer’s Institute, University of South Florida, USA.

### 8.2 Alzheimer’s Disease Metabolomics Consortium

Andrew Saykin^73^, Kwangsik Nho^73^, Mitchel Kling^13^, John Toledo^13^, Leslie Shaw^13^, John Trojanowski^13^, Lindsay Farrer^51^, Gabi Kastsenmü ller^74^, Matthias Arnold^74^, David Wishart^75^, Peter Wü rtz^76^, Sudeepa Bhattcharyya^77^, Cornelia van Duijin^78^, Lara Mangravite^79^, Xianlin Han^80^, Thomas Hankemeier^73^, Oliver Fiehn^82^, Dinesh Barupal^82^, Ines Thiele^83^, Almut Heinken^83^, Peter Meikle^84^, Nathan Price^84^, Cory Funk^84^,Wei Jia^86^, Alexandra Kueider-Paisley^30^, P. Murali Doraiswamy^30^, Jessica Tenebaum^30^, Colette Black^30^, Arthur Moseley^30^, Will Thompson^30^, Siam Mahmoudiandehkorki^30^, Rebecca Baillie^30^, Kathleen Welsh-Bohmer^30^, Brenda Plassman^30^.

73 Indiana University

74 Helmholtz Zentrum Muenchen

75 The Metabolomics Innovation Centre, Canada (TMIC)

76 Nightingale Health

77 University of Arkansas

78 Erasmus MC

79 SAGE Networks

80 University of Texas Health Science Center, San Antonio

81 Leiden University Metabolomics Center

82 West Cost Metabolomics Center

83 University of Luxembourg

84 Baker Heart and Diabetes Institute

85 Institute for Systems Biology

86 University of Hawaii

