## Supplementary Materials for "A blood-based signature of cerebrospinal fluid A*β*_1–42_ status" (from another version of this preprint: version 2, 2018-03-28)

### A blood-based signature of cerebral spinal fluid $A\beta_{1-42}$ status

[1]IBM Australia - Research, Carlton, Victoria, Australia
[4]School of Psychological Sciences, University of Melbourne, Victoria, Australia
[2]Centre for Epidemiology and Biostatistics, The University of Melbourne, Parkville, Australia
[3]Department of Computing and Information System, The University of Melbourne, Parkville, Australia
[4]A complete listing of ADNI investigators can be found at: http://adni.loni.usc.edu/wp-content/uploads/how_to_apply/ADNI_Acknowledgement_List.pdf.
[5]A complete listing of ADMC investigators can be found at:
https://sites.duke.edu/adnimetab/about-us/the-team/
[6]The Florey Institute for Neuroscience and Mental Health, The University of Melbourne, Parkville, Australia
[*]These authors contributed equally to this work

#### ABSTRACT

It is increasingly recognized that Alzheimer's disease (AD) exists before dementia is present and that shifts in amyloid beta occur long before clinical symptoms can be detected. Early detection of these molecular changes is a key aspect for the success of interventions aimed at slowing down rates of cognitive decline. Recent evidence indicates that of the two established methods for measuring amyloid, a decrease in cerebral spinal fluid (CSF) amyloid $\beta_{1-42}$ ($A\beta_{1-42}$) may be an earlier indicator of Alzheimer's disease risk than measures of amyloid obtained from Positron Emission Topography (PET). However, CSF collection is highly invasive and expensive. In contrast, blood collection is routinely performed, minimally invasive and cheap. In this work, we develop a blood-based signature that can provide a cheap and minimally invasive estimation of an individual's CSF amyloid status using a machine learning approach. We show that a Random Forest model derived from plasma analytes can accurately predict subjects as having abnormal (low) CSF $A\beta_{1-42}$ levels indicative of AD risk (0.84 AUC, 0.78 sensitivity, and 0.73 specificity). Refinement of the modeling indicates that only *APOEε4* carrier status and four analytes are required to achieve a high level of accuracy. Furthermore, we show across an independent validation cohort that individuals with predicted abnormal CSF $A\beta_{1-42}$ levels transitioned to an AD diagnosis over 120 months significantly faster than those predicted with normal CSF $A\beta_{1-42}$ levels and that the resulting model also performs reasonably across PET $A\beta_{1-42}$ status.
This is the first study to show that a machine learning approach, using plasma protein levels, age and *APOEε4* carrier status, is able to predict CSF $A\beta_{1-42}$ status, the earliest risk indicator for AD, with high accuracy.

[†]Data used in preparation of this article were generated by the Alzheimer's Disease Metabolomics Consortium (ADMC). As such, the investigators within the ADMC provided data but did not participate in analysis or writing of this report.

[‡]Data used in preparation of this article were obtained from the Alzheimer's Disease Neuroimaging Initiative (ADNI) database (adni.loni.usc.edu). As such, the investigators within the ADNI contributed to the design and implementation of ADNI and/or provided data but did not participate in analysis or writing of this report.

### 1 Supplementary methods

#### 1.1 Download date

The data used in this study were obtained from the Alzheimer's Disease Neuroimaging Initiative (ADNI) database, http://adni.loni.usc.edu/, downloaded $10^{th}$ October 2017, as the R package, ADNIMERGE.

#### 1.2 Plasma protein level measurements

Plasma samples were analyzed by Rules-Based Medicine (RBM) (Austin, TX) using the Human DiscoveryMAP multiplex immunoassay[1]. This immunoassay panel of 190 analytes (Table 'rbmqc' in ADNIMERGE) included proteins previously reported to be involved in cell-signaling and/or associated with a variety of disease processes, including AD, metabolic disorders, inflammation, cancer, and cardiovascular disease. Of the 190 analytes, 146 met previously described quality control procedures (details at http://adni.loni.usc.edu/wp-content/uploads/2010/11/BC_Plasma_Proteomics_Data_Primer.pdf). We also examined total plasma homocysteine (Table 'hcres' in ADNIMERGE), $A\beta_{1-40}$ and $A\beta_{1-42}$ levels in plasma (Table 'upennplasma' in ADNIMERGE) and neurofilament light chain (NFL, Table 'blennowcsfnfl' in ADNIMERGE) for each patient, given these have previously been associated with increased Alzheimer's disease risk.

#### 1.3 Plasma metabolite level measurements

Metabolites from plasma samples were measured using the AbsoluteIDQ-p180 kit, with an ultra-performance liquid chromatography system. This targeted metabolomics approach led to measures of 186 endogenous metabolites (Tables 'admcdukep180fia' and 'admcdukep180uplc' from ADNIMERGE) quantitatively (amino acids and biogenic amines) and semiquantitatively (acylcarnitines, sphingomyelins, PCs, and lyso-glycero-phosphatidylcholines across multiple classes). A multi-stage process was developed by The Alzheimer's Disease Metabolomics Consortium (ADMC) to remove missing data, duplicated and generally poorly measure metabolites, resulting in 138 metabolites that pass this stringent QC[2]. In this work, we made use of the QC'd tables provided by ADMC in the supplementary methods of their related publication[2].

#### 1.4 CSF A$\beta_{1-42}$ measurements

CSF A$\beta_{1-42}$ was measured using a Luminex xMAP platform and Innogenetics INNO-BIA AlzBio3 immunoassay (Table 'upennbiomk2' in ADNIMERGE), as described previously[3].

#### 1.5 Data preparation

Any variables that had more than 15% missingness were removed from the analysis. The only exception was NFL, which was only present in those individuals that also had corresponding CSF measures. The pattern of missingness for remaining variables was checked across the three diagnosis groups and no clear trend could be observed. No individual had more than 5% of variables marked as missing. All remaining missing data points were imputed using an unsupervised RF approach[4].

#### 1.6 Univariate analysis

For the baseline analysis (Table 1 in the main paper), we evaluated the univariate association of demographic variables, *APOE*$\varepsilon$4 carrier status and CSF A$\beta_{1-42}$ levels with diagnosis as well as normal, abnormal CSF A$\beta_{1-42}$ categories. For continuous variables, analysis of covariance (ANCOVA, type II) analysis with post hoc linear hypothesis testing (Tukey comparison between groups) was performed. Normality of variables was checked by visual inspection of the histograms. Box-Cox analysis of the variables was also performed to find the most appropriate power transformation. For CSF A$\beta_{1-42}$, a square root transformation was required. For the analyses of the CSF measurements, *APOE*$\varepsilon$4 carrier status was included as a covariate. For categorical variables, chi-squared tests were performed.

#### 1.7 Random Forest

Random Forests (RF) were selected as the modeling approach to predict CSF A$\beta_{1-42}$ levels, for both the binary and regression tasks. RFs are a widely used ensemble method that builds many simple classifiers (decision trees) in the training phase, each constructed over different sample "bags", a set of samples randomly selected with replacement, and different subsets of features. After training, predictions for unseen samples are made by averaging over all individual trees, taking either the mean or the mode for regression or classification tasks respectively. RFs have a number of advantages for the small sample size and disparate types of features observed in the ADNI dataset. RFs are invariant to the scale of the observed features and make few assumptions about the distributions of observed data allowing them to be applied to multiple data modalities easily.

##### 1.8 Metrics used to determine regression cutoff

Six criteria from the the R package *OptimalCutoffs*[9] were used to determine the optimal threshold for the continuous regression predictions. These criteria were:

- MaxProdSpSe - maximizes the product of sensitivity and specificity.
- NPVEqualPPV - finds the threshold whereby the negative and positive predictive value are closest.
- ValueSe - optimizes for a Sensitivity of 0.85.
- ValueSp - optimizes for a Specificity of 0.85.
- MinPvalue - minimizes the p-value from a Chi-squared test between the actual and predicted markers.
- PrevalenceMatching - matches prevalence estimates from actual and predicted labels.

These methods were selected as they represent the three classes of methodologies in the *OptimalCutoffs* package and were intuitive choices. Further choices were restricted due to concerns of runtime and overfitting.

##### 1.9 All R packages used

All analysis was performed with R (3.3.2)[5], and used the following packages:

- RandomForestSRC[6]
- ggplot2[7]
- pROC[8]
- OptimalCutpoints[9]
- Survminer[10]
- Mixtools[11]
- Stargazer[12]
- ggExtra[13]
- stringr[14]
- dplyr[15]
- tidyr[16]
- pbapply[17]
- pander[18]
- data.table[19]
- gridExtra[20]
- caret[21]

##### 1.10 Cohort Breakdown

| Diagnosis | CN | MCI | AD |
|---|---|---|---|
| **N** | 58 | 396 | 112 |
| **Mean age (sd)** | 75.11 (5.77) | 74.74 (7.40) | 74.80 (8.08) |
| **N Females (%)** | 28 (48.28) | 140 (35.35) | 47 (41.96) |
| **Mean years education (sd)** | 15.65 (3.04) | 15.64 (3.04) | 15.09 (3.20) |
| **N *APOEε4* carriers (%)** | 5 (8.62) | 211 (53.28) | 76 (67.86) |
| **N with CSF** | 58 | 198 | 102 |
| **Mean CSF A$\beta_{1-42}$ (sd)** | 250.84 (21.08) | 163.44 (52.76) | 142.69 (39.15) |
| **Mean CSF tTAU (sd)** | 63.62 (21.76) | 102.33 (59.63) | 120.94 (56.51) |
| **Mean CSF pTAU (sd)** | 21.07 (8.43) | 36.11 (19.27) | 42.07 (19.87) |

**Table 1.** Demographic characteristics of the ADNI data used, without stratification by whether CSF A$\beta_{1-42}$ measures were available.

| CSF A$\beta_{1-42}$ category | Normal | Abnormal |
|---|---|---|
| **N** | 119 | 239 |
| **Mean age (sd)** | 75.27 (7.22) | 74.32 (7.40) |
| **N Females (%)** | 44 (36.97) | 92 (38.49) |
| **Mean years education (sd)** | 15.75 (2.96) | 15.52 (3.10) |
| **N *APOEε4* carriers (%)** | 17 (14.29) | 165 (69.04) |

**Table 2.** Demographic characteristics of the training sub-cohort, stratified into CSF A$\beta_{1-42}$ Normal/Abnormal using a threshold of $\leq$ 192 pg/mL for abnormal.

| Diagnosis | CN | | MCI | | AD | |
|---|---|---|---|---|---|---|
| **CSF A$\beta_{1-42}$ category** | **Normal** | **Abnormal** | **Normal** | **Abnormal** | **Normal** | **Abnormal** |
| **N** | 58 | 0 | 53 | 145 | 8 | 94 |
| **Mean age (sd)** | 75.11 (5.77) | - | 74.55 (8.31) | 74.28 (7.18) | 81.04 (7.31) | 74.38 (7.77) |
| **N Female (%)** | 28 (48.28) | - | 12 (22.64) | 53 (36.55) | 4 (50.00) | 39 (41.94) |
| **Mean Years education (sd)** | 15.67 (2.78) | - | 15.98 (3.06) | 15.75 (2.97) | 14.75 (3.69) | 15.16 (3.27) |
| **N *APOEε4* carriers (%)** | 5 (8.62) | - | 12 (22.64) | 94 (64.83) | 0 (0.00) | 71 (75.53) |

**Table 3.** Demographic characteristics of the training sub-cohort, stratified by both baseline diagnosis and CSF A$\beta_{1-42}$ Normal/Abnormal using a threshold of $\leq$ 192 pg/mL for abnormal.

#### 2 Supplementary Results

##### 2.1 Univariate analysis confirms no significant differences across diagnostic groupings

The cohort used in this study consisted of 58 cognitively normal (CN), 396 mild cognitively impaired individuals (MCI), and 112 subjects with Alzheimer's Disease (AD), with a mean follow up time of 4.1 years (2.9 SD). Table 1 in the main text presents the overall characteristics of the cohort, stratified by diagnosis as well as by subjects which have had CSF $A\beta_{1-42}$ measurements (training set) and those who did not (validation set). Univariate analysis across the diagnosis groupings (Supplementary Methods) confirms their similarity. There was no difference in age between the diagnostic groups ($p = 0.978$), nor was there a gender bias ($p = 0.104$). In line with previous studies, AD and MCI subjects had a greater number of *APOEε4* carriers, than CN ($p = 7.52 \times 10^{-13}, p = 4.97 \times 10^{-10}$, respectively). Further, there was no difference in the years of education between the diagnostic groups ($p = 0.219$). The AD subjects had lower CSF $A\beta_{1-42}$ compared to CN and MCI ( $p < 2.221^{-16}, p < 2.221^{-16}$, respectively) subjects. These similarities were also observed within both the training and validation sets. However, across these two sets of individuals, there was a strong difference in the proportions of the diagnostic class, with the validation set containing 198 MCI but no CN and only 10 AD.

#### 2.2 Performance of different feature subsets using only CN and MCI individuals

| Feature set | Regression | | | | | Binary | | | |
|---|---|---|---|---|---|---|---|---|---|
| | AUC | $R^2$ | Acc | Sens | Spec | AUC | Acc | Sens | Spec |
| BPM | **0.805 (0.07)** | 0.241 (0.11) | **0.738 (0.07)** | **0.739 (0.13)** | 0.732 (0.14) | **0.800 (0.07)** | **0.719 (0.07)** | **0.853 (0.09)** | 0.543 (0.15) |
| BM | 0.769 (0.09) | 0.174 (0.13) | 0.709 (0.07) | 0.691 (0.14) | 0.736 (0.14) | 0.766 (0.08) | 0.691 (0.08) | **0.841 (0.09)** | 0.499 (0.15) |
| BP | **0.813 (0.08)** | **0.258 (0.12)** | **0.728 (0.08)** | **0.719 (0.13)** | 0.735 (0.16) | **0.808 (0.07)** | **0.726 (0.08)** | **0.856 (0.09)** | 0.552 (0.17) |
| B | 0.759 (0.09) | 0.229 (0.15) | **0.738 (0.08)** | 0.690 (0.13) | **0.796 (0.15)** | 0.765 (0.09) | **0.714 (0.08)** | 0.698 (0.13) | **0.736 (0.16)** |
| $BP_{fs}$ | 0.787 (0.09) | 0.224 (0.16) | **0.724 (0.08)** | **0.735 (0.13)** | 0.704 (0.15) | | | | |

**Table 4.** Mean and standard deviation (in parentheses) of performance metrics (area under the receiver operator curve, AUC; accuracy, Acc; sensitivity, Sens; specificity, Spec and $R^2$ for the regression models) using individuals that were CN or MCI at baseline, for the different Random Forest models using different feature sets across all cross-validation folds. Left and right halves are for the Regression and Binary tasks respectively. Bold faced text on AUCs indicates the best performing model or those that are statistically equivalent (via a Wilcox rank signed test, with a Bonferroni-corrected significance threshold of 0.05/5=0.01). Features sets describe combinations of (B) baseline model (age and *APOEε*4 carrier status), (P) Proteomics, (M) Metabolomics.

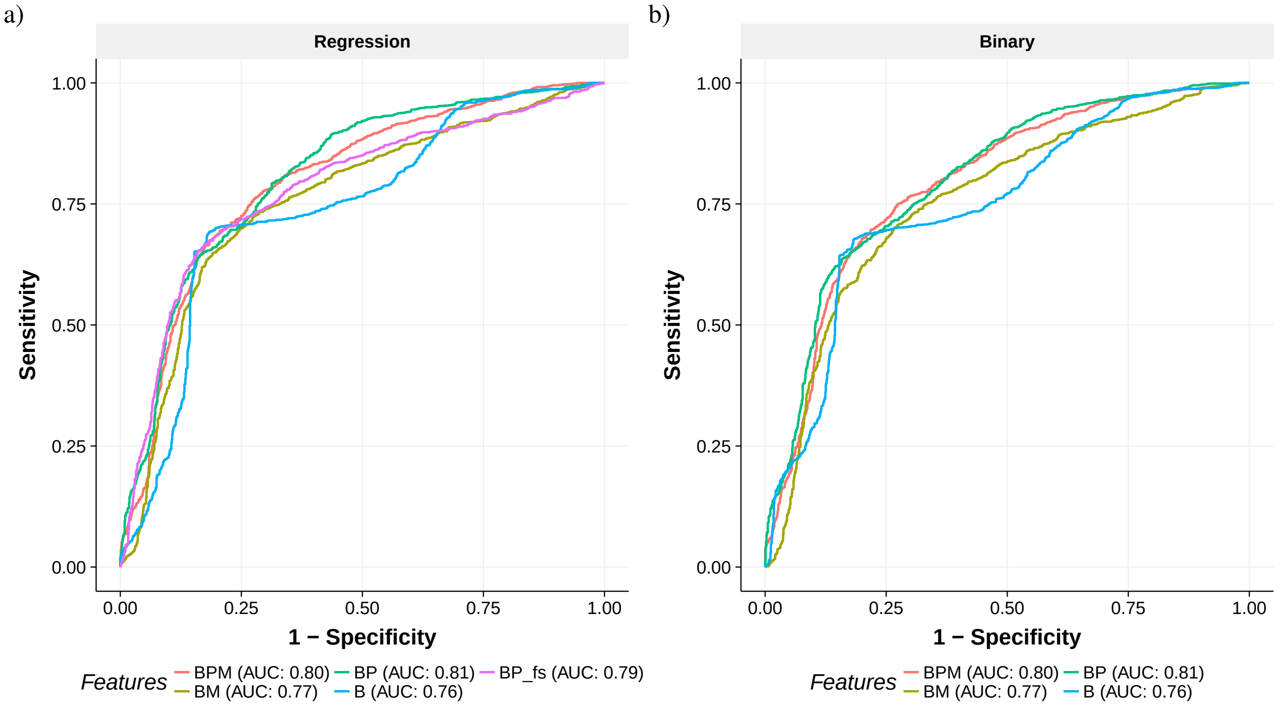


**Figure 1.** ROC curves derived on only CN and MCI individuals in the training cohort, comparing different sets of features to determine predictive value. a) regression, and b) binary. Different colours of lines correspond to different feature sets with corresponding AUCs indicating in the legend under each plot. Feature sets describe combinations of (B) baseline model (age and *APOEε*4 carrier status), (P) Proteomics, (M) Metabolomics.

##### 2.3 ROC curves for different feature subsets using only complete cases

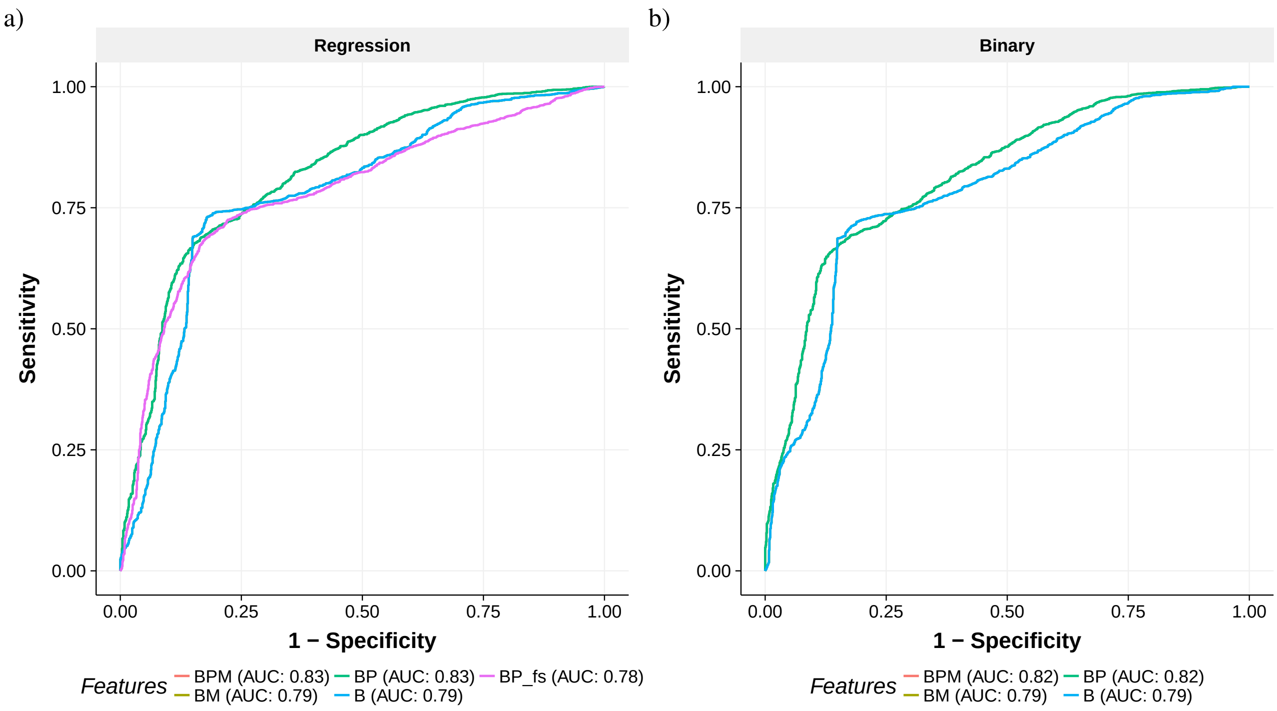


**Figure 2.** ROC curves for a) regression, and b) binary tasks comparing different sets of features to determine predictive value built on 162 plasma analytes with less than 1% missingness, rather than 15% in the main analysis, and 344 samples with no missing analyte measures. Hence this analysis shows the impact of imputation. As the results are very similar to the main analysis, albeit with metabolites have less impact as many were removed, and hence indicates the imputation used has minimal impact on our analysis.
